# Genome-wide translating mRNA analysis following ketamine reveals novel targets for antidepressant treatment

**DOI:** 10.1101/254904

**Authors:** Oliver H. Miller, Nils Grabole, Isabelle Wells, Benjamin J. Hall

## Abstract

Low-dose ketamine is an efficacious antidepressant for treatment-resistant unipolar and bipolar depressed patients. Major Depression Disorder patients receiving a single infusion report elevated mood within two hours, and ketamine’s antidepressant effects have been observed as long as seven days post-treatment. In light of this remarkable observation, efforts have been undertaken to “reverse-translate” ketamine’s effects to understand its mechanism of action. Major advances have been achieved in understanding the molecular, cellular, and circuit level changes that are initiated by low-dose ketamine. Although enhancement of protein synthesis clearly plays a role, the field lacks a comprehensive understanding of the protein synthesis program initiated after ketamine treatment. Here, using ribosome-bound mRNA footprinting and deep sequencing (RiboSeq), we uncover a genome-wide set of actively translated mRNAs (the translatome) in medial prefrontal cortex after an acute antidepressant-like dose of ketamine. Gene Ontology analysis confirmed that initiation of protein synthesis is a defining feature of antidepressant-dose ketamine in mice and Gene Set Enrichment Analysis points to a role for GPCR signaling, metabolism, vascularization, and structural plasticity in ketamine’s effects. One gene, VIPR2, whose protein product VPAC2 acts as a GPCR for the neuropeptide vasoactive intestinal peptide, was characterized in cortex and identified as a potential novel target for antidepressant action.

## Introduction

Depression is the leading psychiatric cause of disability globally^1^, and first-line therapies targeting the monoamine system are ineffective in a significant population of depressed patients^2-4^. In patients that do respond to first-line therapies, weeks to months of continuous treatment are necessary to reach peak therapeutic efficacy. A single infusion of the n-methyl d-aspartate (NMDA) glutamate receptor antagonist ketamine elicits an antidepressant response within 110 minutes of administration^5^. While ketamine has a serum half-life on the order of hours, the antidepressant effects of a single low dose can persist for as long as a week in treatment-resistant MDD patients^5^. What then is the basis for this long-lasting effect? Several groups have reported that ketamine and specific antagonists of GluN2B-containing NMDA receptors rapidly induce protein synthesis: in these studies, differential regulation of a handpicked subset of candidate proteins was observed^6-8^. Targeted western blot experiments, while undoubtedly informative, provide insight only given an a priori hypothesis. Alternatively, other groups have sought out an unbiased understanding of ketamine’s effects on the transcriptional level^9,10^. The results of these studies provide valuable insight into the differential regulation of mRNA expression and gene networks, yet because transcriptional and translational processes are not perfectly coupled, there remains potential for false positives and false negatives when trying to define the mechanism of ketamine’s actions. For example, post-transcriptional regulation of mRNA by RNA binding proteins such as FMRP and ZPB1 acts as a checkpoint between mRNA expression and protein synthesis, and can be dynamically directed by intrinsic and extrinsic cues to suppress, or permit, translation of a given transcript^11-13^. Additionally, NMDAR modulation is known to both promote and repress gene transcription and activate complex transcriptional networks where clear predictions on protein expression changes are difficult.^14^

For these reasons, we sought to assess the complement of mRNAs that are actively translated in response to ketamine, in order to identify the cellular signaling pathways underlying this long-lasting antidepressant effect. Ribosome-bound mRNA profiling involves isolation and deep sequencing of mRNAs being translated at the ribosome at a given point in time^15^, and provides a quantitative and genome-wide assessment of genes differentially regulated in a given condition. Here, we used ribosomae-bound mRNA profiling (RiboSeq) to compare translational regulation in the medial prefrontal cortex (mPFC) of mice injected with either saline, an antidepressant (low) dose of ketamine, or a non-antidepressant (high) dose of ketamine. Gene-set analysis demonstrated that low, but not high dose ketamine strongly upregulated pathways involved in the initiation and regulation of protein synthesis, and revealed significant enrichment of 17 diverse gene sets, and down regulation of 7. We selected VPAC2, a receptor for endogenous vasoactive intestinal peptide (VIP), for further analysis based on large fold-change, high significance, and pharmacological tractability. We demonstrate that VPAC2 is functionally expressed specifically in somatostatin-positive (SOM+) inhibitory interneurons, and that activation of VPAC2 enhances excitability of SOM+ interneurons in mPFC. VPAC2 might serve as a target for future antidepressant interventions and as a target for modulating the SOM+ inhibitory neuron network in mPFC.

## Results

### Calibration of ketamine’s antidepressant effects

Although ketamine’s long-lasting antidepressant-like effects in rodents are robust and replicable, the exact dosage necessary to elicit such an effect varies depending on species, strain, age, sex, and even animal housing and handling conditions^16-20^. Therefore, we first sought to calibrate our in-house antidepressant dose before undertaking genomic analysis (Figure 1A). Compared to saline-injected controls, mice injected with 3mg/kg ketamine displayed antidepressant-like behavior in the forced swim test, 24 hours after a single treatment (Figure 1B). No other dose (10, 50, or 100mg/kg) proved to significantly change immobility time under these conditions, and locomotion rates were normalized before test time (Figure 1B, C). Based on these observations, we selected our experimental groups as saline (control), 3mg/kg (antidepressant), and 100mg/kg (non-antidepressant) for ribosome-bound mRNA profiling (Figure 2).

**Figure 1.**
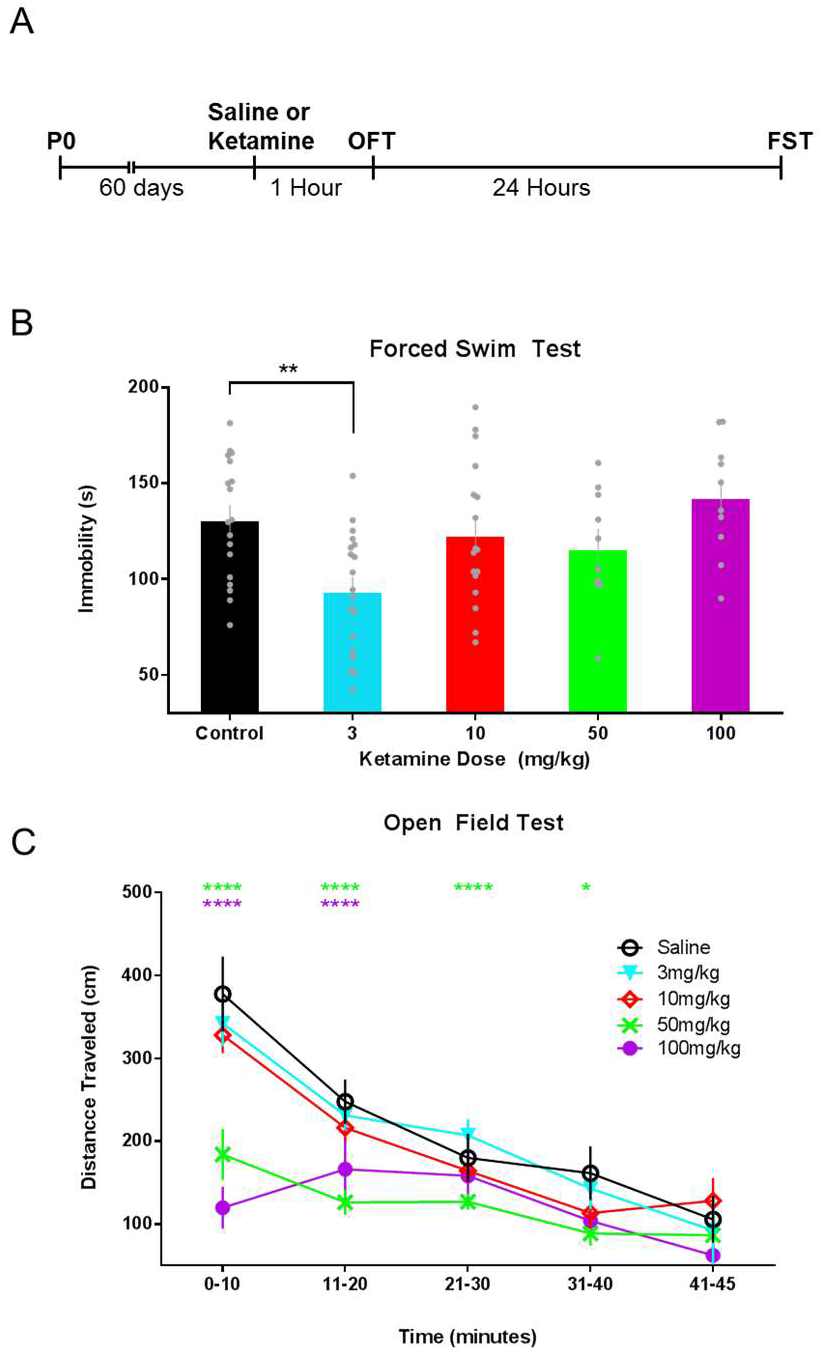
Calibration of ketamine as an antidepressant. (A) Experimental timeline: P60 mice were injected with a single dose of saline or ketamine (3, 10, 50, or 100 mg/kg), and tested in the open field (OFT) one hour later. 24 hours after treatment, depression-related behavior was assessed in the forced swim test (FST). (B) Compared to saline control condition, mice injected with 3 mg/kg ketamine displayed reduced immobility in the forced swim test 24 hours after treatment. No other dose resulted in an alteration in immobility time. (Saline: 131.1±7.43, n=18; Ketamine (3): 93.8±7.22, n=19; Ketamine (10): 123.0±8.56, n=17; Ketamine (50): 116.2±9.69, n=10; Ketamine (100): 142.6±9.72, n=10) One way ANOVA F(4,69) = 4.838, p=0.002, followed by Bonferroni’s multiple comparison test. (C) Antidepressant-like dose ketamine did not alter locomotion of 3mg/kg treated mice. Compared to saline control, mice injected with 50mg/kg and 100mg/kg displayed reduced locomotion in the open field test, which normalized by test end. Two-way ANOVA Treatment: F(4,36) = 4.912, p=0.003), followed by Dunnett’s multiple comparison test. (*p<0.05, **p<0.01, ***p<0.001, ****p<0.0001)

### Isolation of actively translating mRNAs and quality control

Animals from all groups were handled for 5 minutes per day for 7 days before treatment in order to habituate the mice and reduce experimentally-irrelevant translation changes due to handling. Thirty minutes after saline or ketamine treatment, mice were rapidly sacrificed and tissue from mPFC was isolated. Samples were processed to obtain both total mRNAs (for RNA-Seq) and ribosome-bound mRNA fragments (for RiboSeq). Quality control of the data confirmed the prefrontal-cortical identity of the mRNA samples^21^ (Supplemental Figure 1). Bioinformatic analysis revealed a set of differentially expressed genes (DEGs) and differentially translated genes (DTGs) in response to low- and high-dose ketamine treatment.

### Profiling the translatome after antidepressant ketamine

One proposed mechanism by which low-dose ketamine might exert its antidepressant effects is by blocking NMDARs on inhibitory interneurons in mPFC and causing cortical disinhibition^22-24^. Based upon this, one could hypothesize that RNA-Seq analysis would show increases in early immediate gene (IEG) expression in the 3mg/kg ketamine treated mPFC samples. To the contrary, analysis of mRNA abundance showed that 100mg/kg treated mice had increased expression of IEGs Jun, Klf2, Fos, and Klf4, but no significant increase in expression of IEGs were observed in the low-dose (3mg/kg) group (Supplemental Figure 2). This is consistent with other mechanisms of induction including engagement of homeostatic plasticity^24,25^.

**Figure 2.**
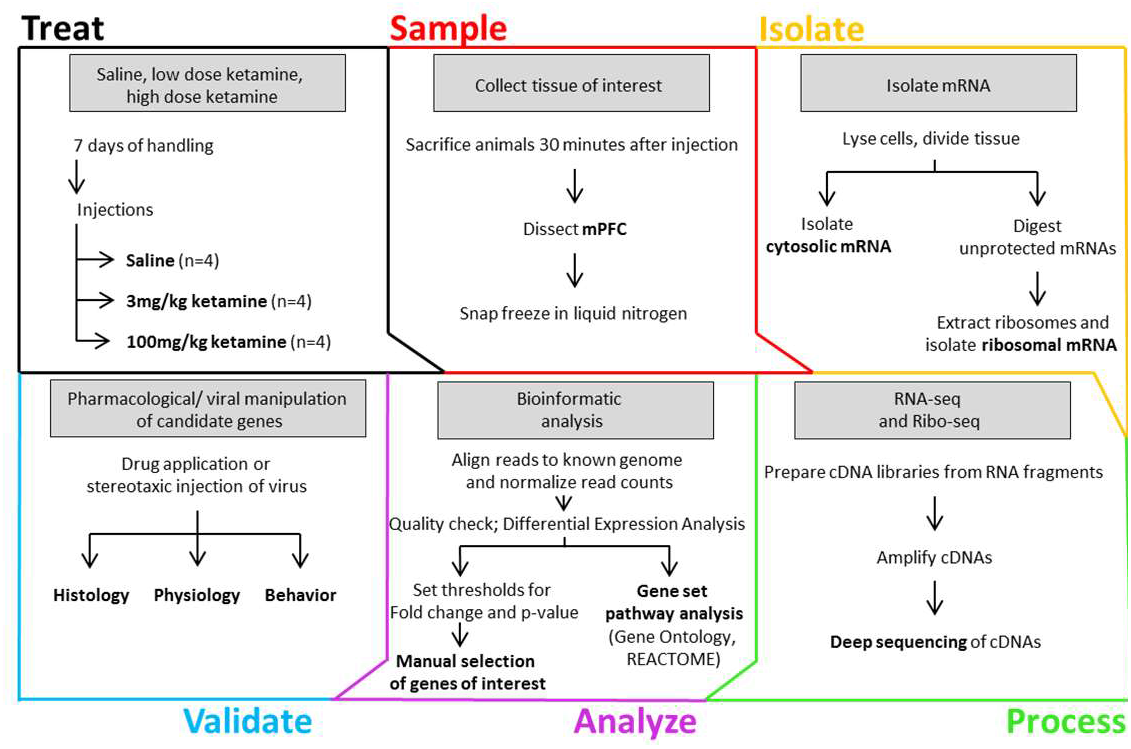
Ribosome-bound mRNA profiling workflow. After seven days of handling, mice were injected with saline, low dose (3mg/kg), or high dose (100mg/kg) ketamine and sacrificed 30 minutes later. mPFC tissue was collected and rapidly frozen. Tissue was processed to obtain total mRNA (for RNA-seq) and ribosome protected mRNA (for ribosome-bound mRNA profiling), and cDNA libraries were produced and sequenced. cDNA reads were mapped to the mouse genome to assess differentially expressed genes and for gene set pathway analysis. Finally, we manually selected candidate genes based on high fold-change, tractability, presumed safety, and novelty.

Dysregulation of biological pathways and functional gene sets contribute to manifestation of depression-related behaviors^26^. Gene Ontology (GO) gene sets are curated based on the functional roles of the contained genes^27^. Analysis of our RiboSeq data set showed that low-dose ketamine treatment drove a robust enhancement of several GO classes relating to the initiation and regulation of protein synthesis including Eukaryotic Translation Elongation, Translation Initiation Complex Formation, and Peptide Chain Elongation (Figure 3A). These results fit well into the existing body of literature, in that initiation of mTOR-dependent protein synthesis is an essential component of ketamine’s long-lasting antidepressant effects^8,28,29^.

**Figure 3.**
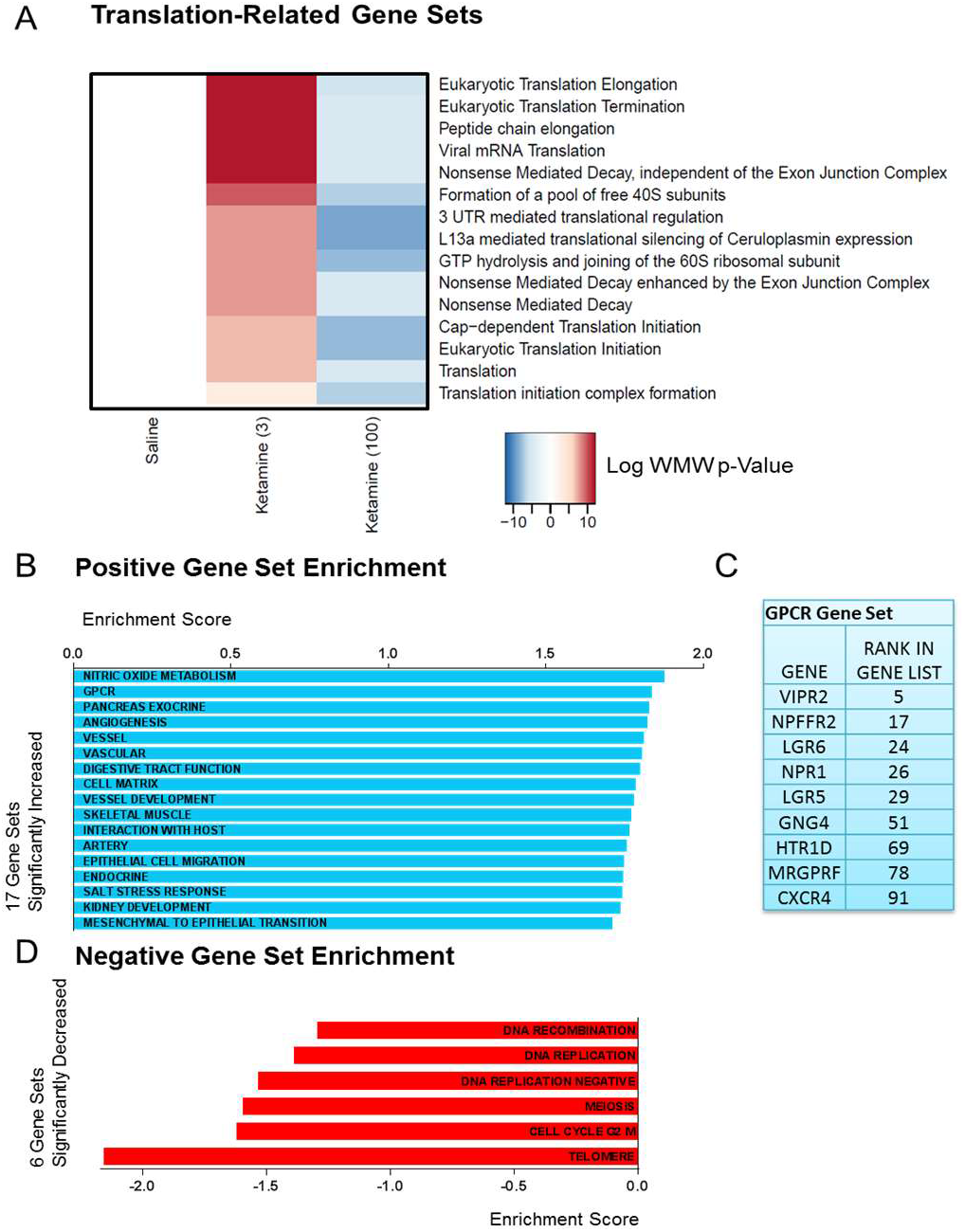
Gene Set Analysis. (A) Gene Ontology (GO) gene set analysis reveals that low-, but not high-dose ketamine treatment results in upregulation of signaling pathways involving translational regulation. (B) Gene Set Enrichment Analysis (GSEA) demonstrates that low-dose ketamine enhances 17 diverse gene sets involved in GPCR signaling, neuronal metabolism, vascularization, and structural plasticity. (C) GSEA gene set “GPCR” enhancement is driven by Vipr2. (D) Six GSEA gene sets are significantly decreased following low-dose ketamine.

In contrast to GO, classification by Gene Set Enrichment Analysis (GSEA) is designed to detect changes in groups of functionally-related genes whose expression is coordinately regulated^30,31^. GSEA analysis revealed that low-dose ketamine alters the expression of diverse gene sets involved in GPCR signaling, neuronal metabolism, vascularization, and structural plasticity (Figure 3B). Of particular interest is the robust 1.83-fold increase in enrichment of the GPCR gene set. This gene set’s enrichment is driven by core enrichment of G-protein GNG4, serotonin receptor HTR1D, and VIP receptor VIPR2, among others (Figure 3C).

Interestingly, the GSEA gene sets that were downregulated after low-dose ketamine are primarily involved in the replication of mitotic cells (Figure 3D). Together, these data suggest that an antidepressant dose of ketamine increases GPCR signaling, metabolism, and structural reorganization, while potentially limiting the genesis of new glia or other mitotic cell types.

### Identification of VPAC2 as a relevant differentially translated gene (DTG) after acute ketamine

Since enrichment analysis pointed to a significant change in genes related to neuronal function such as GPCRs, we set out to identify candidate genes that when differentially translated after low-dose ketamine treatment could drive the antidepressant phenotype. The low-dose ketamine samples contained actively translating mRNAs representing 13,003 total genes (Figure 4A, top). Compared to saline-treated control, 3mg/kg ketamine-treated mice displayed a significant increase in the translation of 61 genes, and a decrease in translation of 22 genes (Figure 4A, bottom).

**Figure 4.**
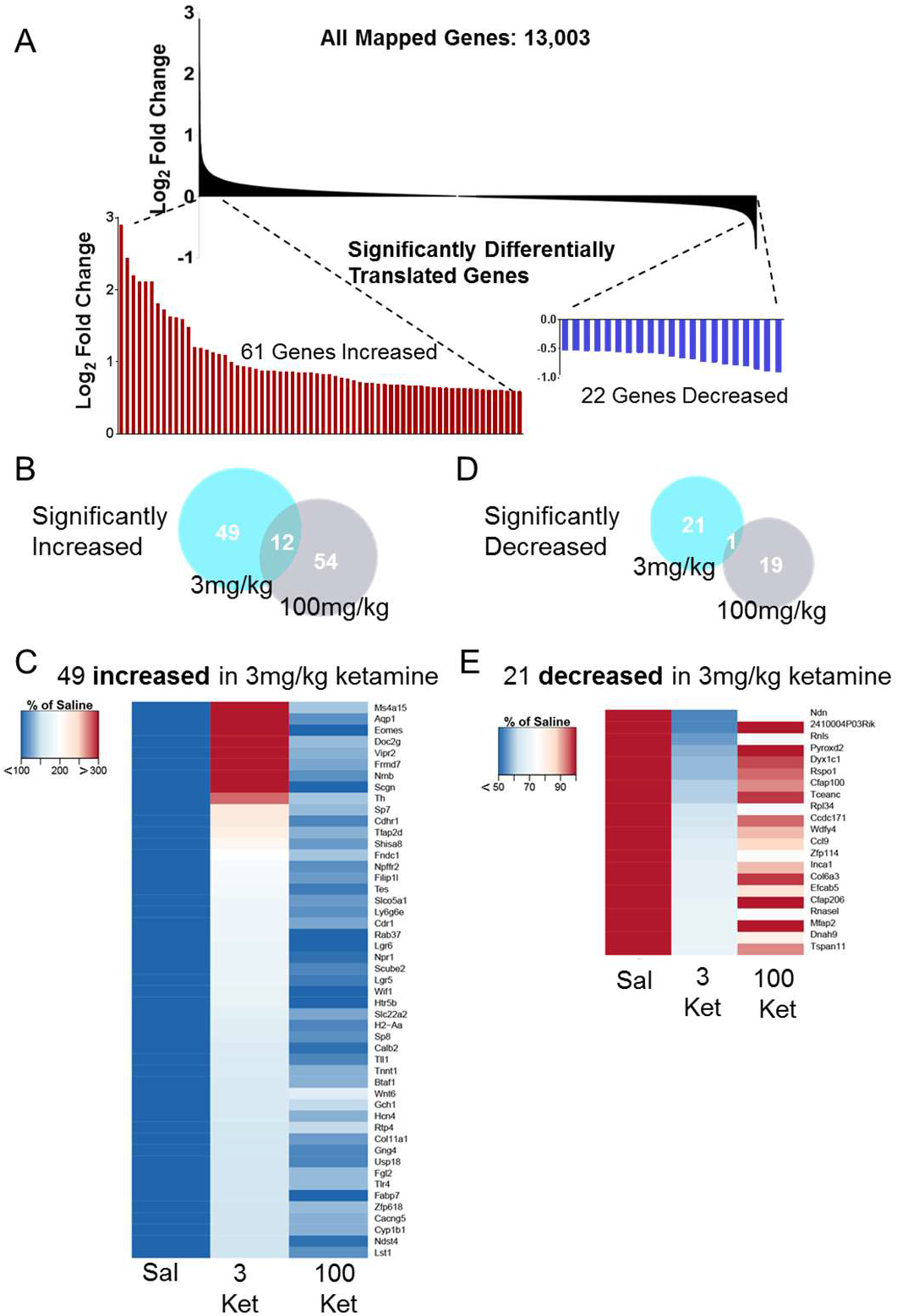
Differentially translated genes after low-dose ketamine. (A) 13,003 ribosome-bound mRNAs were mapped to the genome in the 3mg/kg treatment group. 61 DTGs were increased to greater than 150% of saline control levels, 22 DTGs were deceased to less than 70% of saline control levels. (B) Of the 61 DTGs increased in the 3mg/kg treatment group, 12 were also increased in the 100mg/kg group. These 12 DTGs were excluded from subsequent consideration. (C) 49 DTGs were uniquely increased after 3mg/kg ketamine, normalized to saline control and sorted by percent change in 3mg/kg. Lookup table range limited to 100%-300% of saline expression. (D) Of the 22 DTGs decreased in the 3mg/kg treatment group, 1 was also decreased in the 100mg/kg group. This DTG was excluded from subsequent consideration. (E) 21 DTGs were uniquely decreased after 3mg/kg ketamine, normalized to saline control and sorted by percent change in 3mg/kg. Lookup table range limited to 100%-50% of saline expression.

To ensure that the genes being translated are related to antidepressant-like-dose ketamine, and not due to non-specific NMDAR antagonism-related processes, we subtracted out any genes that were also significantly differentially regulated in mice injected with 100mg/kg ketamine (Figure 4 B, E), and were left with 49 significantly increased DTGs, and 21 significantly decreased DTGs (Figure 4C, F). With this increased confidence in the relevance of the remaining DTGs, we selected target proteins to study further based on the following criteria: The target (1) is pharmacologically tractable, (2) has a favorable prediction for side effects based on previously published data (tissue expression pattern and known phenotypes from knockout or knock-in models), and (3) has no established role in depression. Examples of significantly altered DTGs that were not investigated further because they did not meet these criteria include Ms4a15, Aqp1 and Th, respectively.

Based on the GSEA data pointing to the relevance of GPCRs in antidepressant-dose ketamine’s effects, its high fold-change in the low-dose ketamine treated mice, and satisfaction of our criteria, a prime candidate emerged in VIPR2. The VIPR2 mRNA codes for VPAC2 protein, a membrane bound, Gs-coupled GPCR. Its primary endogenous ligand is VIP, a 28 amino acid neuropeptide synthesized in brain by VIP+ inhibitory interneurons. In the suprachiasmatic nucleus, ultrastructural data suggests that VIP peptide is co-released with GABA from VIP+ interneuron synapses^32,33^, and high frequency stimulation of VIP+ interneurons in the myenteric plexus elicits VIP peptide release and a slow depolarization of connected cells^34^. If this mechanism is conserved in mPFC, the upregulation in expression of VPAC2 we observed after low-dose ketamine treatment would promote a delayed, prolonged excitability in the postsynaptic partners of VIP+ interneurons.

### Role of VPAC2 in the cortical microcircuit

First, we sought to determine which cell types in mPFC functionally express VPAC2. To accomplish this, we crossed a Cre-dependent tdTomato reporter mouse line^35^ to lines expressing Cre in subpopulations of inhibitory interneurons, and patched fluorescent neurons in mPFC. Analysis of spiking parameters in somatostatin positive (SOM+), parvalbumin positive (PV+), and pyramidal neurons (PYR) (Supplemental Figure 3) show expected electrophysiological characteristics. PV+ interneurons displayed a short latency to spike (Supplemental Figure 3B, D) and a shorter action potential half-width (Supplemental Figure 3C, D) compared to SOM+ and PYR neurons. Both PV+ and SOM+ inhibitory interneurons displayed lower capacitance compared to PYRs (Supplemental Figure 3F), and SOM+ cells showed higher membrane resistance relative to PV+ or PYR neurons (Supplemental Figure 3G).

VIP interneurons in cortex have recently become the focus of an intense effort of study^36-40^. In the cortical microcircuit, VIP+ interneurons preferentially inhibit (SOM+) interneurons via fast GABA_A_R-mediated neurotransmission. Given the established cytoarchitecture, and the published expression patterns in cortical neurons^41^, we hypothesized that SOM+ neurons might express VPAC2 and be VIP peptide-sensitive.

To test this hypothesis, we assessed the effects of VPAC2 activation on excitability of these three mPFC neuronal cell types. Under conditions of pharmacologic synaptic blockade (NBQX, APV, Picrotoxin), we injected depolarizing currents sufficient to elicit 5±1 action potentials for SOM+ and PYRs, and the minimum amount of current to elicit consistent firing in PV+s. After 5 minutes of stable, repeated action potential responses, we washed in the selective VPAC2 agonist Bay55-9837 (Bay55), and measured the effects on spiking in these neurons (Figure 5). When recording from SOM+ neurons, perfusion of Bay55 increased the number of action potentials elicited to a maximum of 260% (p<0.01)(Figure 5A). Inclusion of GDPβs in the patch pipette prevented the excitatory effects of Bay55 (Supplemental Figure 4), suggesting that VPAC2 agonism activates SOM+ interneurons by activating G-protein-coupled signaling pathways such as activating adenylyl cyclase and elevating intracellular cAMP. Although VIP+ interneurons preferentially innervate SOM+ interneurons, they also form synaptic contacts with PV+ and PYR neurons^42,43^. In contrast to SOM+ neurons, when recording PV+ or PYR neurons, perfusion of VPAC2 agonist had no significant effect on action potential firing (Figure 5B,C). These results suggest a functionally cell-type specific expression of VPAC2, which when activated leads to preferential excitation of SOM+ interneurons in mPFC.

**Figure 5.**
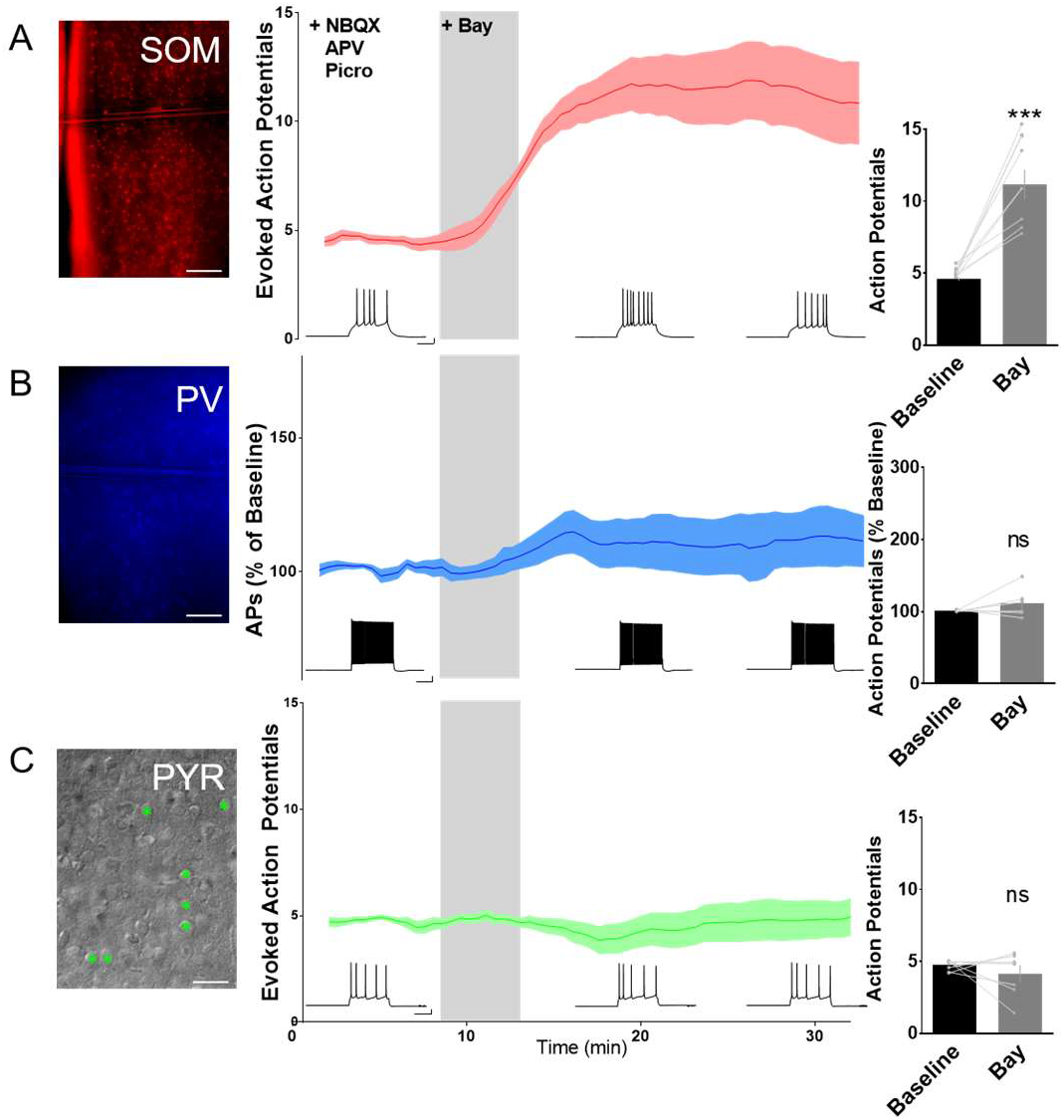
VPAC2 activation enhances action potential firing in SOM+ interneurons in mPFC. effect (A) After establishing a baseline of excitability in response to a stable, repeated current injection, perfusion of VPAC2-specific agonist Bay55 resulted in a significant enhancement of action potential firing in Somatostatin+ interneurons in mPFC. (pre drug: 4.59±0.099 APs, post drug 11.16±1.994 APs, n=9) Paired t-test p=0.0002. (B) Perfusion of VPAC2-specific agonist Bay55 did not alter action potential firing in Parvalbumin+ interneurons in mPFC. (pre drug: 101.3±0.464% of baseline APs, post drug 111.8±8.43% of baseline APs, n=6) Paired t-test p=0.277. (C) Perfusion of VPAC2-specific agonist Bay55 did not alter action potential firing in layer II/III Pyramidal neurons in mPFC. (pre drug: 4.426±0.136 APs, post drug 4.136±0.579 APs, n=7) Paired t-test p=0.371. Visually identified pyramidal neurons in DIC are marked with green asterisks. (*p<0.05, **p<0.01, ***p<0.001, ****p<0.0001)

## Discussion

One hypothesis as to how ketamine exerts its antidepressant-like effects posits that ketamine preferentially antagonizes NMDARs on inhibitory interneurons, thereby disinhibiting pyramidal neurons. Disinhibited pyramidal neurons then tonically fire causing cortical hyperactivation, glutamate release, and LTP-like plasticity. The disinhibition hypothesis proposes that this LTP-like plasticity promotes synaptogenesis in mPFC pyramidal neurons, and an antidepressant-like effect. Disinhibition occurs in deep-layer prelimbic mPFC after NMDAR antagonist MK801 (0.1mg/kg) treatment^23^ and results in elevated levels of PFC glutamate as measured by microdialysis^44^. However, GluN2B-specific antagonists do not elicit cortical disinhibition^45^, yet still elicit an antidepressant-like response, suggesting that cortical disinhibition might be a correlative effect, and not causal for ketamine’s antidepressant-like effects. Accordingly, we found that supra-antidepressant doses, but not antidepressant doses of ketamine drove cortical activation, as measured by the increased transcription of immediate early gene mRNAs 30 minutes after treatment. This data suggests that cortical disinhibition need not occur for ketamine’s antidepressant effect.

Interestingly, GO gene set analysis confirms previous demonstrations that robust initiation of protein synthesis is a hallmark of antidepressant ketamine treatment, and not a nonspecific effect of NMDAR antagonism generally^6,7^. While we found that several GO gene sets related to translational regulation were significantly increased after low-dose ketamine, conversely, these same pathways were downregulated with the high, non-antidepressant-like dose. GSEA analysis points to roles for GPCR signaling, neuronal metabolism, vascularization, and structural plasticity in mPFC underlying ketamine’s long-lasting antidepressant-like effect, both supporting the existing literature^46-54^, and opening up new avenues for further exploration.

The GPCR gene set and the GPCR VPAC2 were of particular interest to us based on high fold change, pharmacological accessibility, and the fact that its role in depression is unstudied. We identified that action potential firing in SOM+ inhibitory interneurons is uniquely enhanced by VPAC2 agonism in mPFC.

The functional specificity of VPAC2 to SOM+ interneurons is an interesting finding in light of recently produced cell-type specific RNA-seq data, which demonstrates Vipr2 mRNA present in not only SOM+, but also in PV+ and Reelin/HTR3A+ interneurons^55^. As VPAC2 is primarily a Gs-coupled GPCR, it is possible that in PV interneurons Gs signaling does not link to pathways that regulate cell excitability. Alternatively, it is possible that cAMP activation does not couple to intracellular pathways that directly alter firing probability in PV interneurons.

Although a role for VPAC2 in major depression has not, to our knowledge, been previously identified, there is precedence for a role in neuropsychiatric disease. Copy number variations in the gene coding for VPAC2, Vipr2, have been identified to confer a significant risk for schizophrenia^56^, pointing to the importance of this gene for maintenance of normal brain function. It would be interesting to see more detailed genotype/phenotype analysis of these scizophrenia patients in terms of the severity of negative symptom endophenotypes such as amotivation.

These findings show promise for the prospect of identifying novel targets for antidepressant treatment based upon the translational profile of known fast-acting antidepressants. Identifying these cell or circuit-specific molecular handles will allow us to address symptoms in a domain-specific manner, potentially enhancing efficacy and broadening the window between intended effects and undesired side effects.

## Experimental Procedures

### Animal experimentation

This study was performed with strict adherence to the Swiss federal ordinance on animal protection and welfare as well as according to the rules of the Association for Assessment and Accreditation of Laboratory Animal Care International (AAALAC), and with the explicit approval of the local cantonal veterinary authorities. Mice were housed on a 12/12hr light-dark cycle, at ∼20°C and ∼55% humidity. Regular rodent chow and tap water were available ad libitum. Mice were weaned at ∼P30 by gender. Juveniles were group-housed to not more than three mice per cage. Male mice 2-3 months of age were used at the time of analysis for all physiology, behavior, and histology studies

### Mice

#### GluN2B-Floxed Mice

Mice with exon 5 of the GRIN2B gene flanked by loxP sites were used for all experiments to maintain the same background as experiments from previous studies^8,57^. At no point in this study were GluN2B-floxed mice exposed to Cre, and therefore these mice are wild-type at the protein level.

#### Ai14 Mice

The Ai14 mouse line (Rosa-CAG-LSL-tdTomato-WPRE: ΔNeo) has a LoxP-flanked STOP cassette preventing transcription of downstream tdTomato cDNA incorporated into the Rosa locus^58^. Expression of Cre-recombinase results in Cre-LoxP recombination and deletion of the STOP cassette, allows expression of tdTomato in cells of interest. Ai14 mice were crossed with SOM-cre and PV-cre mice for cell identification.

#### Somatostatin Cre Mice

Sst::cre (Jax stock: 013044) mice have a SOM-IRES-Cre knock-in allele with an internal ribosome entry site and Cre recombinase in the 3’ UTR of the somatostatin locus. When SOM-IRES-Cre mice are bred with mice containing loxP-flanked sequences, Cre-mediated recombination will result in deletion of the floxed sequences in the SOM-expressing cells in the offspring.

#### Parvalbumin Cre Mice

Parvalbumin::Cre (Jax stock: 008069)mice have an IRES-cre-pA knock-in allele with an internal ribosome entry site and Cre recombinase in the 3’ UTR of exon 5 of the parvalbumin locus. When PV Cre mice are bred with mice containing loxP-flanked sequences, Cre-mediated recombination will result in deletion of the floxed sequences in the PV-expressing cells in the offspring.

### Library preparation and sequencing

Preparation of ribosome-bound mRNA profiling and total RNA sequencing libraries using the TruSeq Ribo Profile kit (Illumina, #RPHMR12126) was performed as detailed in the manufacturer’s protocol. In brief, frozen brain tissue samples were ground with a pestle on dry ice and thawed in the presence of lysis buffer containing 1% TritonX-100, 0.1% NP40, 1mM DTT, DNase (0.01 U/μL) and 100 μg/mL Cycloheximide. Three quarters of the debris-cleared lysate was used for ribosome footprinting by digestion with RNAse for 45 minutes at room temperature. The remaining quarter of the sample was used for RNAseq library prep workflow, followed by isolation of the ribosome protected mRNA fragments by Phenol-Chloroform extraction.

For ribosomal RNA (rRNA) depletion the RiboZero magnetic Gold kit (Illumina) was used. The following steps of PAGE purification, end repair, adapter ligation, reverse transcription and circularization were performed as described in the manufacturer’s protocol, and final libraries were purified once more on a PAGE gel to remove residual primers.

Quality of amplified libraries was assessed by capillary electrophoresis with a high sensitivity DNA chip on a 2100 Bioanalyzer (Agilent Technologies) and quantified by quantitative PCR with a sequencing library quantification kit (KAPA Biosystems) on a Roche Light Cycler 480. Multiplexed libraries with 1 % spiked in PhiX control were sequenced on a HiSeq2500 instrument for 50 cycles using version 4 chemistry reagents (Illumina).

### Bioinformatic analysis

Data processing and bioinformatics analysis was performed as previously described^59^. In short, linker tags were removed from RNA sequencing and ribosome-bound mRNA profiling reads by the FASTX Toolkit, all reads that mapped to rRNAs, tRNAs or mitochondrial rRNAs were removed, and the remaining reads were mapped to RefSeq (v38) by TopHat. Finally all read counts that mapped uniquely to genes were extracted for expression analysis. We applied the edgeR algorithm^60^ for differential gene expression analysis and identified genes with change larger than 150% or less than 70% of saline control levels, and Benjamini-Hochberg adjusted p value less than 0.01 as significantly changed. We applied the camera algorithm^61^ for gene set enrichment analysis with gene sets collected in the Roche internal database RONET. Gene ontology enrichment analysis was performed using the Fisher’s exact test.

### Electrophysiology

#### Slice preparation

As previously descibed^8,62^, for acute slice recordings, adult 2-3 months old mice were anesthetized with isoflurane and decapitated. The brain was rapidly removed and placed into ice-cold modified NMDG solution (composition in mM: 110 NMDG, 110 HCl, 3 KCl, 10 MgCl2 6*H20, 1.1 NaH2PO4 H20, 0.5 CaCl2 dihydrate, 25 glucose, 3 pyruvic acid, 10 ascorbic acid, 25 NaHCO3, 305-310 mOsm). A vibratome (Leica VT1200S) was used to generate 350μm PFC coronal sections containing the mPFC. Slices were allowed to recover in oxygenated (95% oxygen, 5% CO2) NMDG for 30 minutes at 34°C, then transferred to a holding chamber where they were incubated in bicar¬bonate buffered ACSF (composition in mM: 124 NaCl, 4 KCl, 26 NaHCO3, 1.26 NaH2PO4, 3 MgSO4, 2 CaCl2) at room temperature for at least another 30 minutes before whole-cell recordings. ACSF solutions were bubbled with 95%O2/5%CO2 at all times to maintain consistent oxygenation and pH.

#### Whole-cell patch-clamp recordings

For current clamp recordings, borosilicate glass pipettes were filled with a potassium-gluconate intracellular solution (120 K Gluconate, 10 Hepes, 10 D-Sorbitol, 1 MgCl2 6H20, 1 CaCl2, 1 EGTA, 5 Phosphocreatine disodium salt hydrate, 2 ATP disodium salt hydrate, 0.3 GTP sodium salt, pH adjusted to 7.3, 290 mOsm). Whole cell patch clamp recordings were acquired (MultiClamp 700B, Axon Instruments) and digitized sampled at 20kHZ (Digidata 1550, Axon Instruments), filtered at 2kHz and acquired with pClamp software. Clampfit (Axon Instruments), MiniAnalysis (Synaptosoft), and custom-written macros in IgorPRO (Wavemetrics) were used to analyze raw data, and GraphPad Prism was used to perform statistical analyses, as noted in text. Pipette resistances ranged 2-5 MΩ Series access resistance ranged from 4 to 20 MΩ and was monitored for consistency. Cells were discarded if leak current rose above 200 pA, access resistance was greater than 20MΩ or changed by more than 15% during recording. Layer III neurons confined to prelimbic and infralimbic mPFC were targeted for recording, observable with epifluorescent imaging.

At baseline for SOM and PYR neurons, action potentials were evoked by small current injections, sufficient to elicit 4-6 action potentials. For PV neurons, a baseline of 4-6 APs was unattainable, so the minimum stimulus necessary to evoke burst firing was used. After a minimum of 5 minutes of baseline recording, Bay55 ([maximal] 367.1±19.33nM, as assayed by mass spectrometry) was perfused into the bath for 5 minutes, and the recordings continued unadjusted for the entirety of recording (up to one hour). Graphs are displayed as running averages of three sequential sweeps.

### Mouse Behavior

#### Open Field Test

Open field test was conducted using Accuscan open field system. Mice were introduced into the arena at a fixed corner of an acrylic (42 × 42 × 30.5 cm) field arena and allowed to explore the arena freely. The arena was cleaned in between subjects with 70% ethanol.

#### Forced Swim Test

Forced swim test was carried out as previously described^63,64^. Briefly, mice were introduced to a cylinder filled with room temperature water. The cylinder was 15 cm in diameter by 20cm height and filled to 15 cm (sufficient to prevent mice from using tails to support themselves) with room temperature water (23-26°C). Two to four mice were tested simultaneously with a screen to separate the cylinders. Videos were recorded from a top-mounted camera for 6 min. After the testing, mice were rubbed dry and placed under an infrared lamp for about 20 min. Immobility in the last 4 min of 6 min test session was scored blind to genotype and treatment. The cylinders were cleaned in between subjects with 70% ethanol.

## Author Contributions

O.H.M and B.J.H designed all experiments and wrote the manuscript. N.G. performed library preparation and sequencing steps. I.W. performed the bioinformatic analysis. O.H.M performed all other experiments and analyses.

## Acknowledgments and disclosures

We would like to thank Jitao David Zhang for generating the ribosome-bound mRNA profiling analysis pipeline, and Nikolaos Berntenis for assistance with bioinformatics analysis.

O.M., N.G., I.W., and B.J.H were employees of F. Hoffmann-La Roche Ltd., Basel, Switzerland at the time of experimentation.

## Supplemental Figures

**Supplemental Figure 1.**
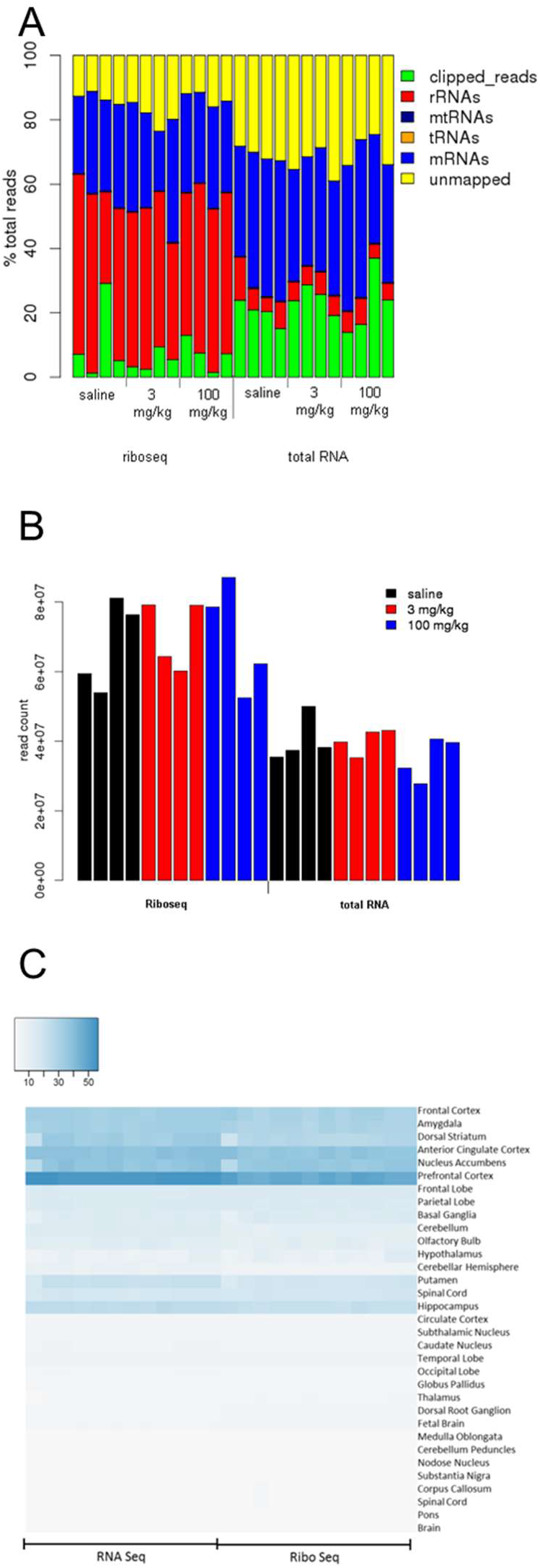
Quality control. (A) RNA-seq and ribosome-bound mRNAs distributed into clipped reads, ribosomal RNAs (rRNAs), mitochondrial RNAs (mtRNAs), transfer RNAs (tRNAs), messenger RNAs (mRNAs), and RNA reads that could not be mapped to the genome. (B) Read counts of mRNAs are suitable for RNA-seq and ribosomal profiling. (C) Both RNA-seq and ribosomal profiling mRNAs show high Prefrontal Cortex character in BioQC analysis, confirming tissue identity.

**Supplemental Figure 2.**
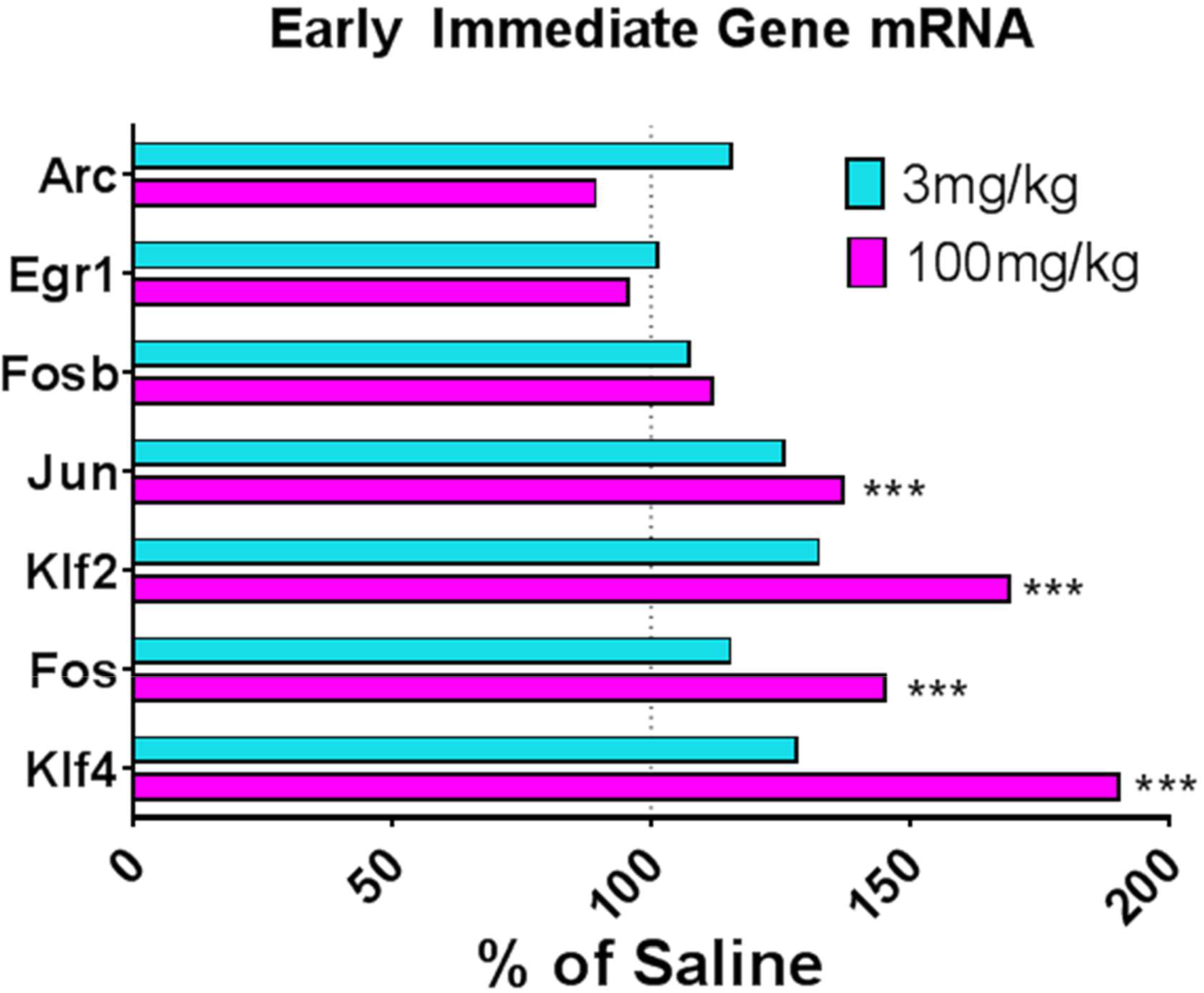
RNA-seq assessment of immediate early genes. RNA-seq assessment of immediate early genes Analysis of immediate early gene mRNA expression demonstrates that high-dose, but not-low dose ketamine initiates activity-dependent expression of mRNAs, relative to saline control (dotted vertical bar). (*p<0.05, **p<0.01, ***p<0.001, ****p<0.0001)

**Supplemental Figure 3.**
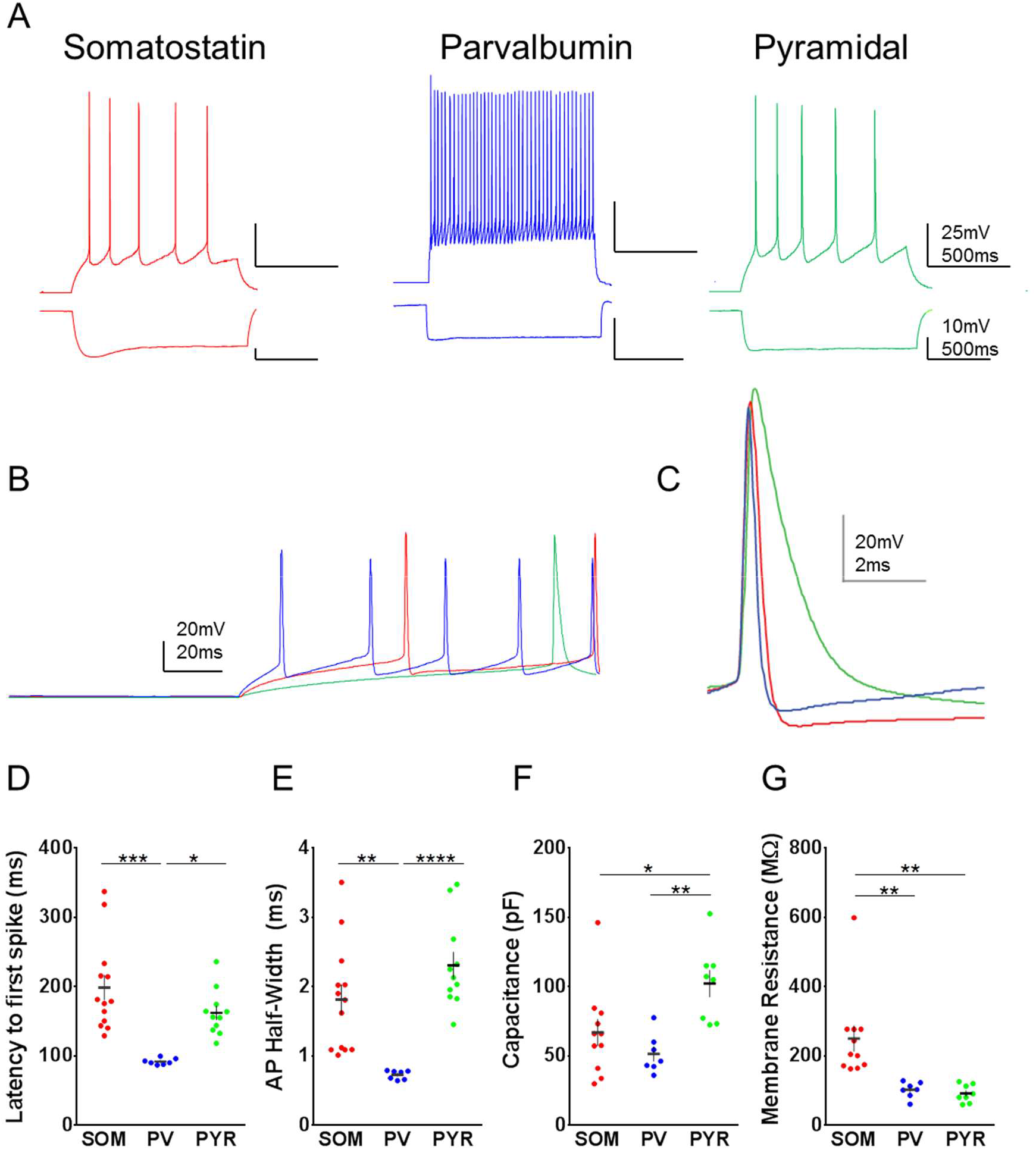
Characterization of cell types in mPFC. (A) Three neuronal subtypes characterized in mPFC: somatostatin+ interneurons, parvalbumin+ interneurons, and excitatory pyramidal neurons. (B) Sample traces showing latency to spike of different cell types, colored as in panel A. (C) Sample traces showing action potential shapes of different cell types, colored as in panel A. (D-G) Different cell types demonstrate unique character of spike latency, action potential duration, capacitance, and membrane resistance. One-way ANOVA with Sidak’s multiple comparisons test. (*p<0.05, **p<0.01, ***p<0.001, ****p<0.0001)

**Supplemental Figure 4.**
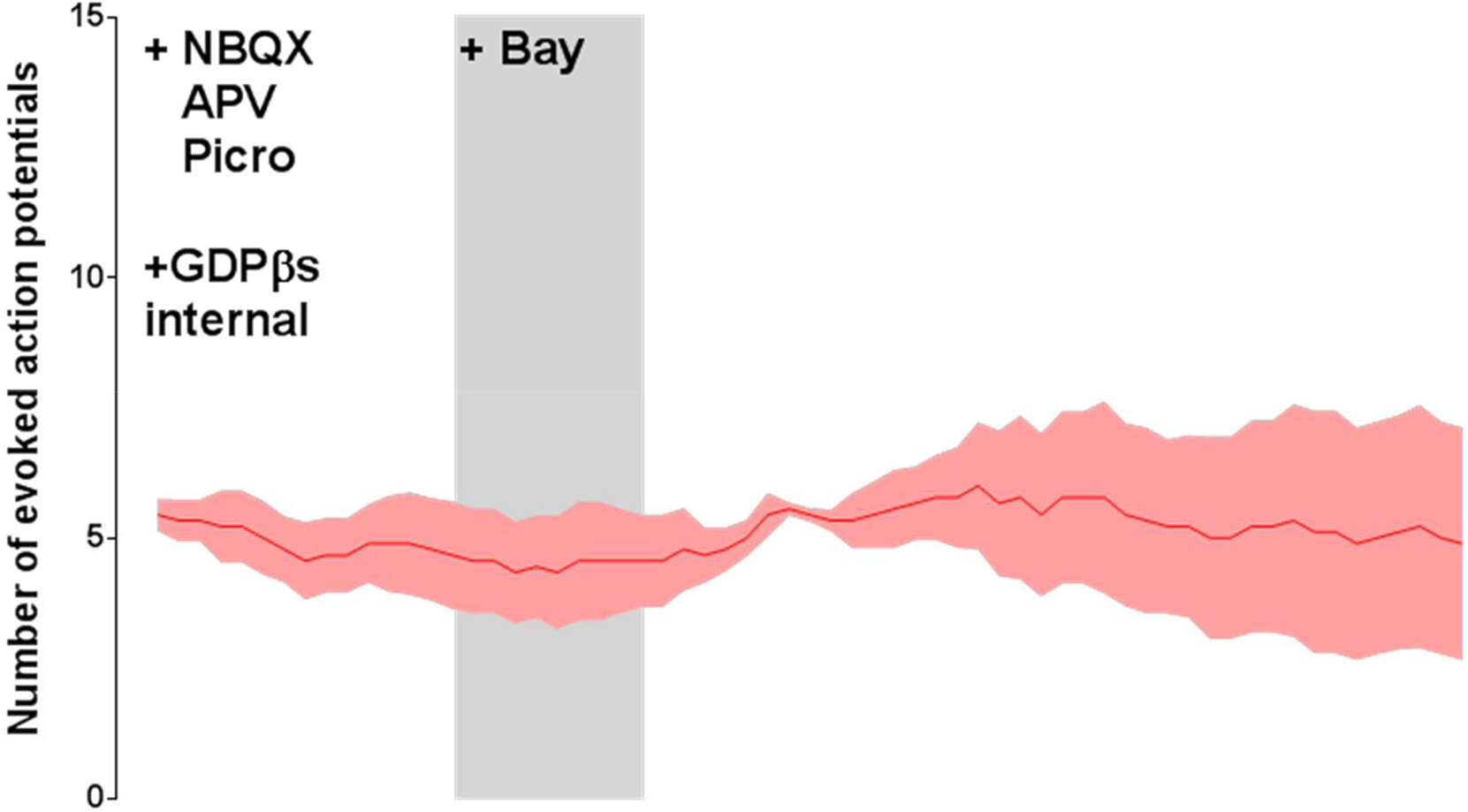
Activation of SOM+ interneurons by VPAC2 agonism is G-protein-dependent. After establishing a baseline of excitability in response to a stable current injection, perfusion of VPAC2-specific agonist Bay55 did not alter action potential firing in Somatostatin+ interneurons in mPFC when GDPβS was included in the patch pipette.

